# Precision-Based Filtering Facilitates Integration of Conventional and Single-Nucleus Transcriptomes to Identify Time- and Temperature-Sensitive Cell Populations

**DOI:** 10.64898/2026.01.28.702137

**Authors:** Adam Seluzicki, Travis A Lee, Nolan T Hartwick, Todd P Michael, Joseph R Ecker, Joanne Chory

## Abstract

Transcriptome analysis via RNA sequencing (RNAseq) has become a ubiquitous method of molecular characterization from whole organisms, dissected tissues, and single cells. These experiments have provided an extraordinary volume of data describing the molecular states and responses to many conditions. However, standard approaches to RNAseq analysis commonly use expression level filters that eliminate potentially useful data in the service of decreasing noise. Here we describe the implementation of a coefficient of variation-based filter for RNAseq gene expression data. This filter prioritizes consistent data across replicates, allowing lowly-expressed genes with low-variation measurements to be retained for downstream analysis. We show that, in our *Arabidopsis* RNAseq data set, this filter allows for the inclusion of many more transcription factors than even a low-stringency expression level filter. We find that these lowly-expressed genes mark specific cell clusters in our single-nucleus (sn)RNAseq dataset. We further characterize communities of co-expressed genes, sampled across the day at two growth temperatures, in relation to snRNAseq cell clusters, finding evidence for a highly photosynthetic cell population, and a cell state marked by high cell division and translation. These methods can be expanded to RNAseq analysis in many systems, facilitating the construction of more detailed models of tissue-specific gene regulatory networks.

## INTRODUCTION

Gene expression profiling is central to understanding many aspects of biology. High-throughput sequencing enables gene expression profiling on a massive scale through RNA sequencing (RNAseq). Analysis of RNAseq data has become routine. Many protocols have been published describing the processing and analysis of the resulting data starting from the raw sequence data, through quality control, alignment, gene-level quantification, normalization, and finally to the determination of differential gene expression between experimental conditions^1–4^. A persistent problem in RNAseq analysis is how to filter out noisy or uninterpretable parts of the data to obtain the most useful information possible. Typically this step is carried out using gene-level count data that has been normalized to the size of the sequencing library for each sample, such as counts per million reads (CPM)^1,4–7^. Most protocols use expression level as the primary criterion for this filter, choosing an expression level below which genes are excluded from downstream analysis. Filtering out lowly expressed genes is often justified by the assumption that low expression is also characterized by noisier measurements and, occasionally, by explicit statements that genes expressed at a low level are, by definition, unlikely to be important^1,5,8^. While it is true that lowly expressed genes are subject to noise, the assumption that the relative importance of a gene is determined by, or correlates with, its expression level is contradicted by many studies of individual genes^9–12^. It is therefore important to refine this filter to retain information on lowly expressed genes that would be discarded under current analysis practices. The recovery of these data, by reanalysis of existing data sets, can potentially provide a wealth of new information and serve to generate hypotheses as to the functions of the many uncharacterized lowly expressed genes.

While most RNAseq experiments are done using bulk RNA extracted from whole organism, organ, or tissue samples, the advent of single cell or single nucleus RNA sequencing (sc/snRNAseq) provides an opportunity to understand the molecular functions within individual cells within a tissue- or organ-level context. However, sc/snRNAseq experiments can be cost-prohibitive, particularly for a large number of samples covering different genotypes, timepoints, or treatment conditions^13^. Recent studies have described high-resolution sc/snRNAseq data sets in a range of plant tissues and developmental stages^14–19^. It is desirable, then, to devise strategies to computationally combine the information in RNAseq and sc/snRNAseq data sets to determine possible cell-type-specific relationships.

In *Arabidopsis* and other species, a large proportion of genes are expressed in responses to the day-night cycle. The circadian clock directly controls some, while others follow environmental stimuli such as light and temperature^20^. While many studies have carried out diurnal and circadian time course gene expression analyses using RNAseq, only one has done so using snRNAseq, sampling plants under free-running conditions in 24h and 48h time courses^21^. This study focused primarily on identifying genes that oscillate with similar patterns in distinct cell types, as well as characterizing the core clock components across cell types. A recent study linked bulk and single-cell RNAseq using a machine-learning tool to estimate the proportions of cell types represented in bulk RNAseq data^22^. This approach uncovered distinct gene expression profiles in subpopulations of leaf mesophyll cells. This study provides proof of principle that coordinated analysis of bulk and single cell RNAseq datasets can yield novel insights. It emphasizes that new analyses using previously published datasets offer a valuable and economical way to derive hypotheses for future testing in the wet lab.

Here, we describe a strategy for RNAseq analysis that replaces the commonly used expression level filter step with a filter based on the coefficient of variation (CV) among replicates. We examine the effects of this filter on the composition of the resulting transcriptome compared with expression level filtering. We find that genes specifically retained by the CV filter are enriched for transcription factors, which would otherwise have been discarded, and that genes with higher noise that are discarded by the CV filter are enriched for genes involved in biotic stress. Using these data, we query for lowly-expressed genes within a snRNAseq dataset, finding restricted expression in specific cell clusters, many of which are of unknown cell type. We note examples of cluster-specific gene expression with highly time- and temperature-dependent expression dynamics, suggesting cell- or tissue-specific environmental responses. Co-expression analysis identified communities of genes with shared expression dynamics across the day. Several of these communities map to specific cell clusters, suggesting specialized temperature-sensitive cell populations or functions, as well as possible transient cell states, rather than stable cell types. Together, these studies describe novel analysis strategies that can provide insight into combinations of large data sets. We use these methods to highlight known genes and processes with uncharacterized time- and temperature-dependent expression, as well as potential cell type- or tissue-specific experimental targets.

## RESULTS

### A strategy for filtering RNAseq data based on the precision of replicates

We sought to examine the information discarded during conventional RNAseq analysis, and to devise a strategy to distinguish between truly unusable data and data that is merely excluded due to expression level. This study focuses on our previously published RNAseq data set, which is both technically and biologically suited to this approach^23^. First, this data set was sampled in biological triplicate, providing a basis for assessing the precision of expression measurements. Second, it was sequenced to an average depth of ∼74 million reads per sample, thereby increasing signal from lowly-expressed transcripts.

Our analysis pipeline (Fig. 1A) uses gene-level counts, such as those from HTseq, as input^24^. These counts are normalized to Counts Per Million Reads (CPM) using the edgeR package^2^. The coefficient of variation (CV) is calculated for each gene across the three biological replicates per condition. In this data set, there are 42 Col-0 samples across 14 conditions (two temperatures, seven time points across the day). CV is calculated on CPM rather than raw reads because the range of read counts between samples before normalization may reflect differences in library size more than technical or biological variation in expression. The mean CV calculated for each gene is used as the filtering criterion, and genes exceeding the threshold are removed. Gene-level counts retained by this filter are re-run through edgeR to normalize to a new library size and arrive at CPM, which is used for downstream analysis.

**Figure 1:**
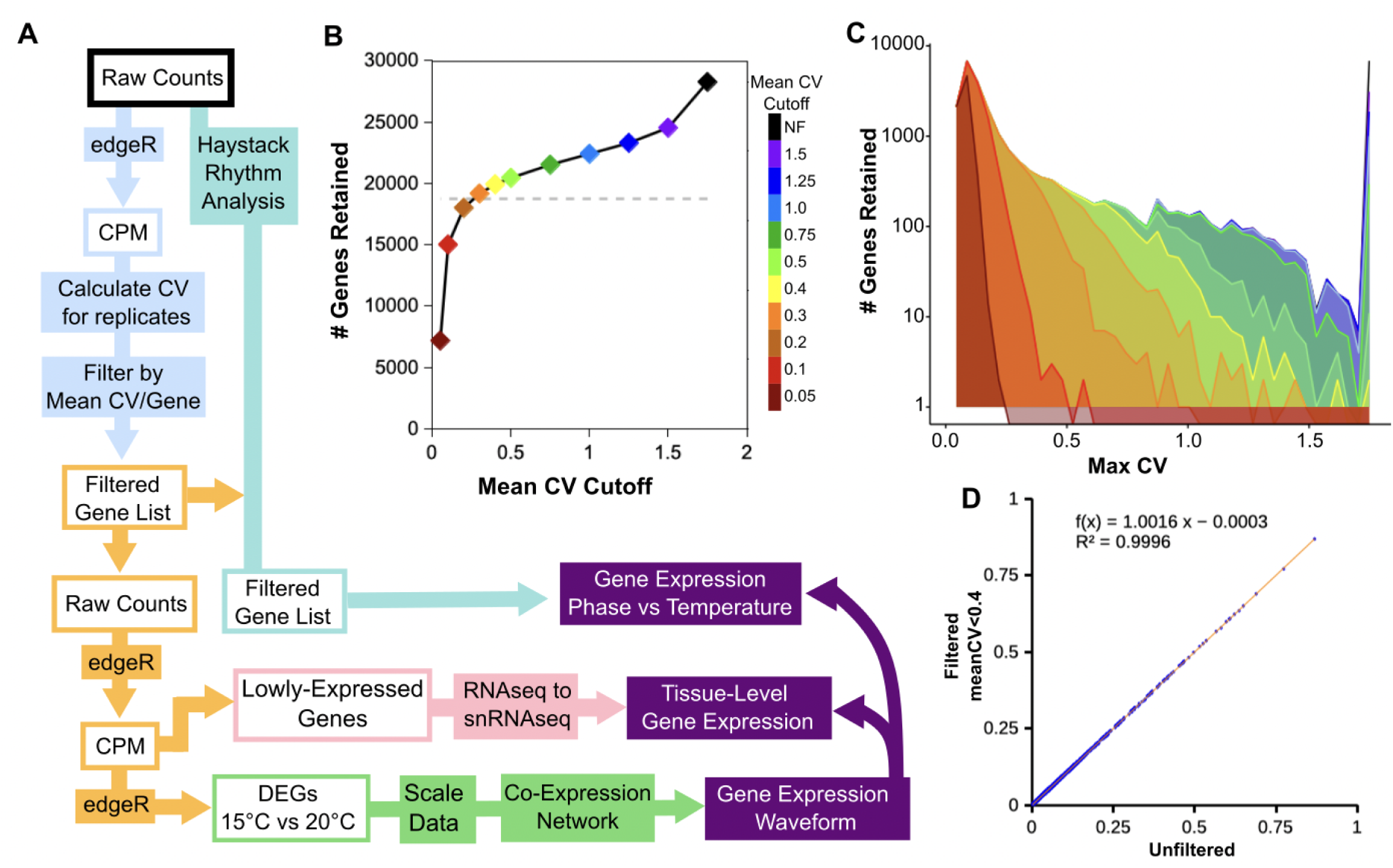
Schematic and quality control for coefficient of variation-based filtering of RNAseq data. (A) Schematic representation of data filtering and analysis workflow. Starting from non-normalized gene-level transcript counts (Counts, black box at top), data are normalized to counts per million reads (CPM) using edgeR (light blue boxes). Coefficient of variation (CV) of experimental replicates is calculated, and the average CV across all conditions is calculated for each gene. Genes with a mean CV below a threshold are retained for downstream analysis. Data for the genes retained by this filter are normalized to CPM, and differential expression is calculated using edgeR (orange boxes). Lowly expressed genes, defined here as genes with average expression below 1 CPM per sample, are selected and mapped to cell clusters defined by snRNAseq (pink boxes). CPM data for differentially expressed genes (DEGs) are scaled to focus on expression dynamics rather than expression level. Pairwise Pearson correlations between genes are calculated and used to build a co-expression network, which defines communities of genes with similar temporal expression dynamics (green boxes). In a parallel analysis pipeline, peak expression phase is calculated using the Haystack algorithm. Gene expression phase differences between plants grown at 20 °C and 15°C are calculated (teal boxes). Co-expressed genes defined by the network analysis are mapped to cell clusters defined by snRNAseq, and examined for temperature-dependent effects on peak phase of expression (purple arrows). (B) The number of genes retained by the meanCV filter at defined thresholds is shown. Grey dashed line indicates the number of genes retained using the expression level filter at >1 CPM. Right side y-axis indicates the percentage of genes retained. Color key at right indicates the meanCV cutoff used in (B) and (C). (C) Maximum CV per gene retained at different meanCV cutoffs. (D) CV of replicates in 15°C at ZT0 for the first 1000 genes starting at the top of Chromosome I present in both meanCV filtered and unfiltered gene lists. Calculated CV is minimally affected by the change in library size after filtering.

We examined the characteristics of this filter, quantifying how many genes are retained at a series of mean CV (mCV) thresholds (Fig. 1B). After an initial rapid decrease(∼14%) between no filter and mCV<1.5, there is a shallow, nearly linear decrease in the number of genes retained from mCV<1.5 (86%) to mCV<0.3(68%), followed by a steep reduction from mCV<0.2 to mCV<0.05 (63% to 25%). The transcriptome size using mCV thresholds between 0.2 and 0.3 is comparable to that of our previously used expression level criterion^23^ (grey dashed line). As this filter is based on the mean CV, it is still possible for genes with large variation between replicates to be retained, provided that low variation is observed in the other conditions. We examined the distribution of maximum CVs of genes retained by the range of mCV thresholds (Fig.1C) and found that the mCV filter progressively reduces the retention of high-maxCV genes. This filter uses data normalized by library size, and the library size changes after the high-variation genes are removed and the data are normalized in the second stage. We tested whether the change in library size may alter the calculated variation among replicates. We examined the CV of the first 1000 genes on Chromosome I that are retained by the mCV<0.4 filter, compared to the CV of those genes in the unfiltered data (Fig. 1D). We found these genes to be nearly identical, with a slope of 1.0016 and R^2^ of 0.9996, suggesting that the change in library size associated with this filter does not meaningfully alter the statistical properties of the replicates.

### Filter type comparison reveals gene type selectivity

The mCV filter retains data on genes that are expressed at low levels provided that the replicates indicate consistent measurement. We examined the effect of each filter type on library size and found that the CPM filter removes an average of ∼46,000 reads per library. In contrast, the mCV filter removes an average of ∼5.2 million reads per sample (Fig.2A-B). This equates to a retention rate of ∼99.94% for the CPM filter, and ∼93% for the mCV filter (Fig. 2C). Filtering at the mCV<0.4 threshold (hereafter noted as mCV) retains a transcriptome of 19920 genes, while the CPM filter retains 18751 genes (Fig. 2D). Thus, relatively few genes with high variability account for a high proportion of total reads. We examined the biological CV which, combined with technical CV, makes up the total CV and is one of the quality control outputs from edgeR, in unfiltered, CPM>1 filtered, and meanCV<0.4 filtered data (Supp. Fig.1)^25^. Perhaps unsurprisingly, we observe much lower total variation using the mCV filter than the CPM filter (Supp. Fig. 1B,C). We asked which genes were specifically retained by each filter and found that, while most retained genes were common between the two filters, the mCV filter retained 1537 genes excluded by the CPM filter, and excluded 368 genes retained by the CPM filter. Together, these data indicate that the CPM filter, which is more permissive in terms of read retention, is more limiting than the mCV filter in terms of gene retention. The small proportion (∼0.06%) of reads rejected by the CPM filter includes data for 1537 genes. Conversely, the ∼7% of reads rejected by the mCV filter correspond to 368 genes, indicating that the filter has the desired effect of retaining information on lowly-expressed genes.

**Figure 2:**
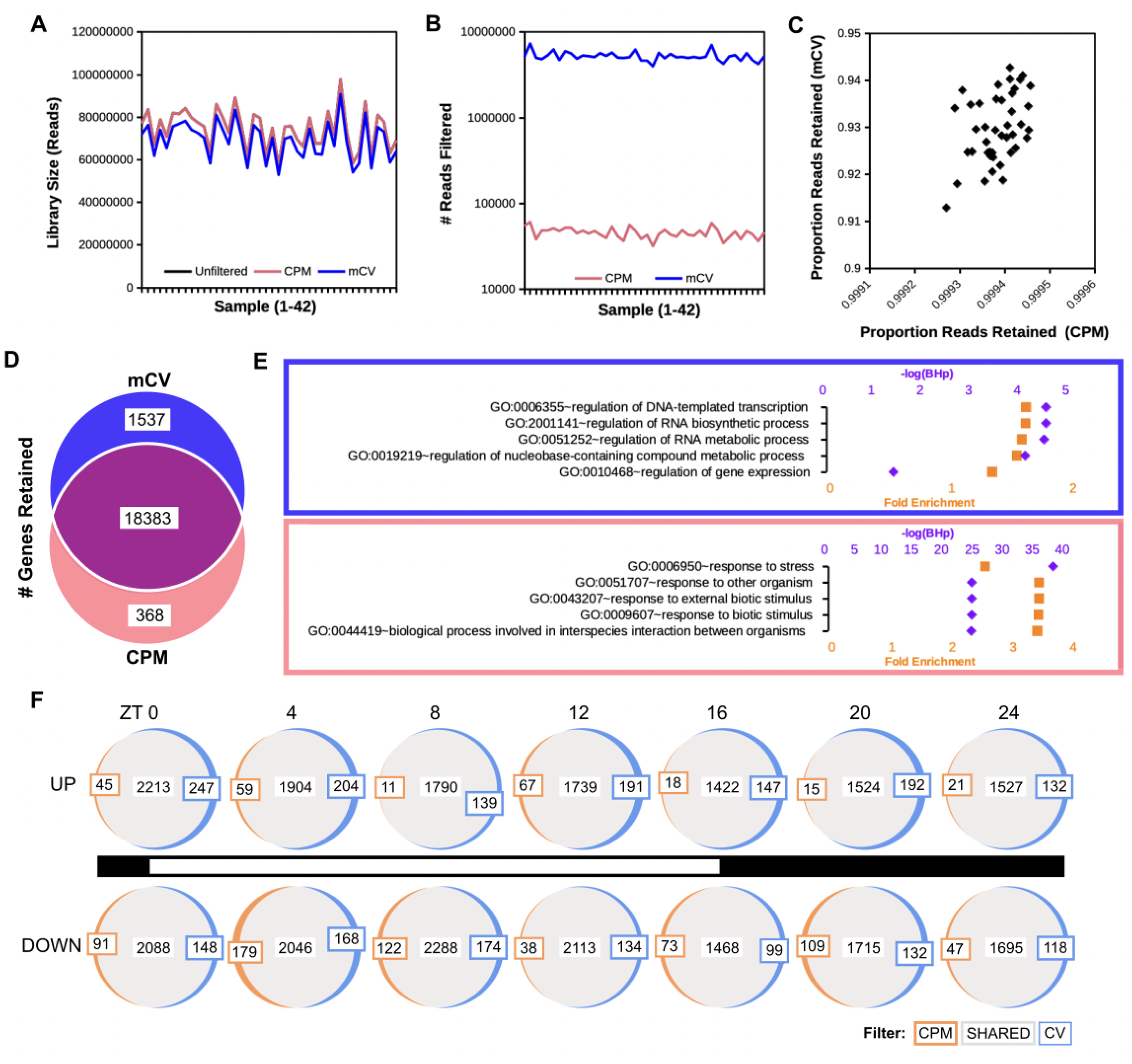
Effects of CV and CPM filters on library and transcriptome composition. (A) Library size across the 42 samples (unfiltered - black, CPM filtered - magenta, mCV filtered - blue). (B) Number of reads removed by filter type (CPM filtered - magenta, mCV filtered - blue). (C) Proportion of reads retained by each filter for each sample. (D) CV filtering selectively retains transcription factors, while the CPM filter retains genes related to biotic stress. A Venn diagram of the number of genes retained using meanCV<0.4 and CPM>1 is shown. Gene Ontology (GO) enrichment was determined for the non-overlapping gene sets. The top five enriched GO categories in each set are shown (blue - retained by CV filter, pink - retained by CPM filter). (E) CV filtering is not selective based on time of day. Venn diagrams show differentially expressed genes identified under meanCV<0.4 (blue) or CPM>1 (orange) filtering, or both (shared - grey) in samples collected across the day. Light conditions are indicated by the black and white bar at center. Sampling time is indicated at top. Lights-on is ZT0, lights-off is at ZT16. Genes identified as up-regulated or down-regulated in 15°C vs 20°C are shown in the upper and lower rows, respectively.

We then asked whether genes specifically retained by each filter were enriched for particular biological processes, and calculated Gene Ontology (GO) enrichment for these subsets (Fig. 2E). The mCV-specific genes were enriched in categories related to transcription, suggesting that this filter retains information about transcription factors. Interestingly, the CPM-specific genes were enriched in categories related to biotic interactions. These data were collected from soil-grown plants rather than plants grown on sterile plate media, and were therefore exposed to a complex biotic environment. This suggests that this filter strategy can identify genes associated with biological experimental variance or genes with inherently variable expression^26^.

We then analyzed the transcriptomes for differential expression between growth temperatures at the seven sampled time points across the day using the mCV- and CPM-filtered data. Consistent with the comparison between the whole transcriptome sizes, differentially expressed genes (DEGs) show more genes included at each time point with the mCV filter than the CPM filter in most cases (Fig. 2F). Genes identified as up- or down-regulated at 15°C vs 20°C show similar trends, and this pattern persists across the day, suggesting it is not time-of-day dependent. Taken together, these data demonstrate that the mCV filter selectively enriches for lowly-expressed genes, including transcription factors.

### Lowly expressed genes map to specific cell clusters identified by snRNAseq

We hypothesized that genes identified as lowly expressed in bulk RNAseq may be either expressed broadly throughout the plant at very low levels, or expressed in specific cell or tissue types. Starting from the mCV transcriptome, we selected the lowest expressing genes (summed CPM across 42 samples of <42 CPM, or average<1 CPM per sample). We examined the expression of these genes within the previously published snRNAseq data set from 12-day-old seedlings, the most similar time point available ^15^. The mean expression of these genes within each of the identified cell clusters, followed by k-means clustering of genes by expression patterns, revealed clear specific expression patterns (Fig. 3A). Of the 19 cell clusters (CC) identified in the previous study, 16 show enriched expression for specific gene clusters (heatmap clusters, HMC). We asked if certain HMCs were more likely to contain DEGs, and found a narrow range of proportions, with no clear association with cell type (Fig. 3B). We found no clear association between the number of DEGs and the number of genes per HMC (Fig. 3C). Focusing on genes that were specifically retained by the mCV filter in each HMC, we found that most contained ∼60% mCV-specific genes (Fig. 3D). We asked if DEGs were over-represented among the mCV-specific genes. Using the full set of mCV-specific genes from the heatmap, we found that 54% were DEGs, compared to an expected 39% (Fig. 3E). We found a similar pattern when DEG proportions in mCV vs mCV+CPM retained genes were calculated for each HMC (Fig. 3F). While one HMC showed a lower-than-expected proportion (31%), most showed statistical enrichment (42-71%) for DEGs among mCV-specific genes.

**Figure 3:**
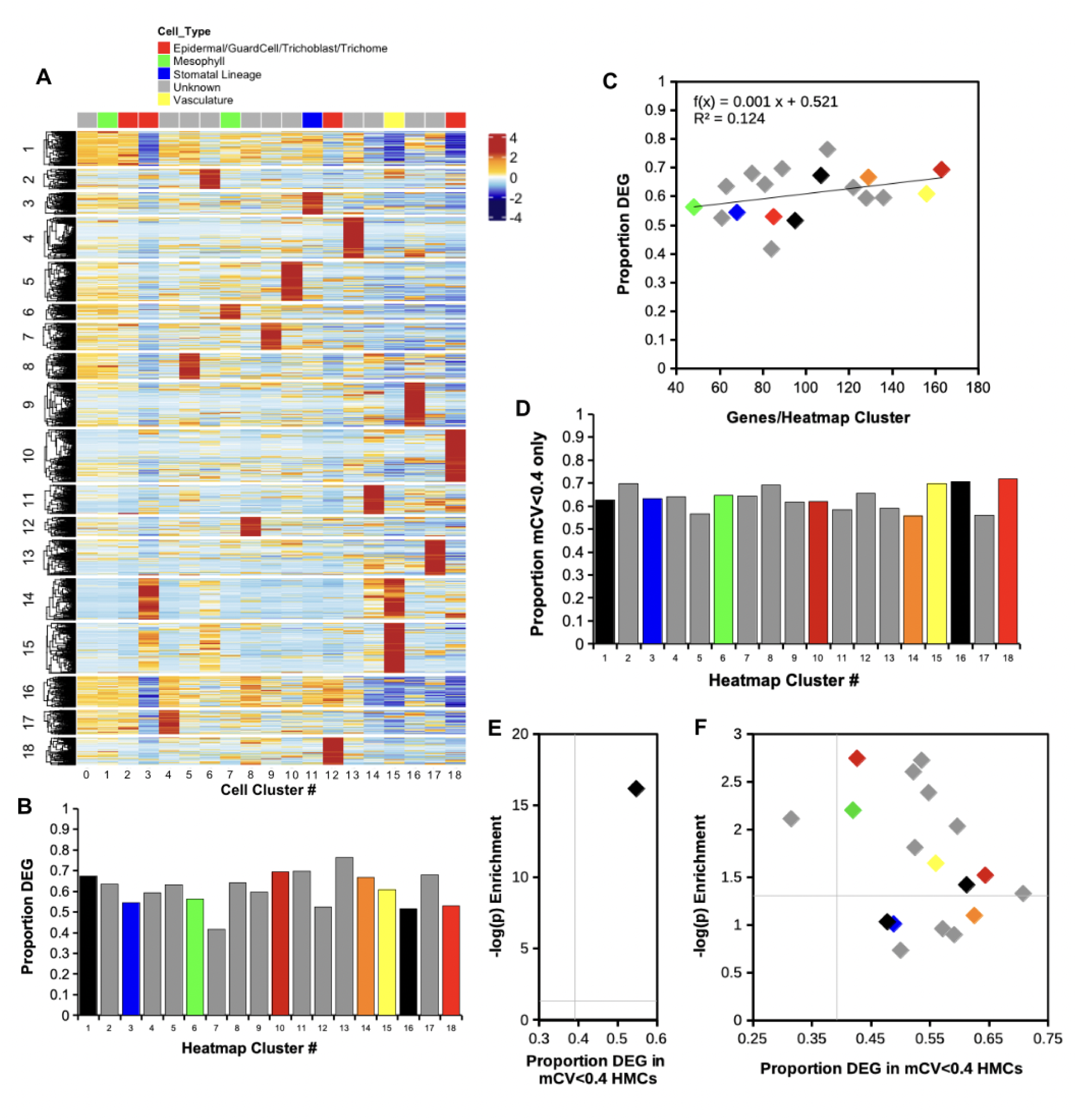
Lowly-expressed genes map to specific cell clusters identified by snRNAseq. (A) Genes retained by the meanCV>0.4 filter with summed expression less than 42 CPM (average expression less than 1 CPM per sample across 42 samples) in time course RNAseq data were selected. Mean expression per cell cluster (0-18) in snRNAseq data are indicated by color, with blue being low expression and red being high expression (color key is at right). Cell clusters are identified by number (see bottom of heatmap) and cell type is indicated by color, with unknown clusters in grey (see Cell_Type key at top). Genes are grouped along the y-axis by k-means clustering. (B) Proportion of DEGs mapping to each gene cluster defined in the y-axis of the heatmap (Heatmap Cluster, HMC). Colors indicate the dominant cell type represented in the cluster as defined at the top of the heatmap. Cluster 14 is enriched for Cell Clusters (CC) 3 and 15 (Epidermal/Trichome and Vasculature, respectively), and is therefore shown in orange. (C) The proportion of DEGS as a function of the number of genes in the HMC. Colors are as in (B). (D) Proportion of genes specifically retained by the mCV filter by HMC. Colors are as in (B). (E, F) Enrichment of DEGs among the genes selectively retained by the mCV filter. (E) Bulk enrichments of DEGs among all mCV-retained genes in this set. (F) Enrichment of DEGS among mCV-retained genes by HMC, colored by CC.

We then mapped the normalized mean expression of HMC genes at the single-cell level. The clustered expression observed in the heatmap, in which we visualized the mean expression of each gene within the cell cluster, persisted when the mean expression of the HMC genes was mapped to individual cells. To learn whether characteristic time- or temperature-dependent expression patterns are embedded in these cell clusters, we examined the expression of these genes in our time course RNAseq data and tested for consistent differential expression. We plotted UMAPs for all HMCs, and the mean expression of all genes in that HMC (Supp. Fig. 2A-R). We found few instances of consistent gene expression dynamics, most of which average to flat scaled expression at 1 across the day. We observed limited statistical differences between temperatures in the combined data (2-way ANOVA + Tukey HSD, Time × Temperature). HMC7, which maps to the uncharacterized cell cluster 9 (CC9), shows higher average expression at 15°C at four time points in the light phase (Supp. Fig. 2G). HMC16, which shows broad expression across cell clusters, shows slightly increased expression at 20°C at four time points (Fig. 3A, Supp. Fig. 2P). Thus, aggregate expression analysis indicates that expression in a cell cluster does not predict temporal gene expression dynamics.

Oscillations in gene expression driven by diurnal cycles and circadian clocks are prevalent throughout biology, and we asked whether the phasing of gene expression responds to temperature, and whether the composition of the CPM and mCV transcriptomes results in differing patterns. We analyzed oscillation parameters for each gene in our RNAseq time-course data using the Haystack algorithm^20^. This algorithm reports phase measurements and confidence estimates: values>0.8 are considered high confidence. We asked if the peak phase of gene expression responds to temperature. We calculated the phase difference for each gene at each temperature. We used the product of the confidence estimates to combine the separate measurements, resulting in a threshold of 0.64. Plotting confidence values against phase differences, we obtained plots showing transcriptome-wide phase difference between the two temperatures under each filter strategy (Supp. Fig. 3A,B). We find that gene expression phase is not strongly impacted by the temperature range tested in this dataset.

We examined six HMC-derived UMAP plots with highlighted cell clusters (Fig. 4). Expression profiles of representative genes specifically retained by the mCV filter and determined to be differentially expressed between temperatures in at least one time point are shown beneath each UMAP. HMC4 maps to CC13 (Fig. 4A). This cluster contains a strongly time- and temperature-regulated, but little-studied DNAJ-domain gene. In plants grown at 15°C, this gene is expressed at ∼0.25 CPM with little change across the day, but expression increases and becomes oscillatory, with a defined peak (∼0.75 CPM) at Zeitgeber Time (ZT)24/0 (lights-on). Expression dips to ∼0.5 CPM from the late morning through early night timepoints. This gene has been shown to regulate the *WUSCHEL* (*WUS*) promoter, and over-expression leads to meristem termination^27^. HMC9 maps to CC16, which is adjacent in the UMAP to CC13 above (Fig. 4B). This cluster contains the NAC-family transcription factor *CUP-SHAPED COTYLEDON 1* (*CUC1*), which is important for the formation of the meristem and the cotyledon organ boundary during development^12,28^. *CUC1* transcript level is temperature-sensitive, with lower expression and mild oscillation peaking in the morning, dropping through the afternoon, and rising again from late evening through the night at 20°C, and elevated and nearly anti-phase expression at 15°C. HMC11 maps to CC14, and contains the transcription factor *HOMEOBOX PROTEIN 52* (*AtHB52*) (Fig. 4C). The *AtHB52* transcript shows peak expression at ZT0, dropping rapidly to a mid-day through evening trough, and increasing again starting at lights-off (ZT16). AtHB52 has been shown to be important for integrating light and nitrogen input to promote turnover of photodamaged photosynthetic apparatus^29^. HMC13 maps to CC17, and expresses the GHMP-kinase family gene *AtGALK2* (Fig. 4D). AtGALK2 may be involved in salt and abscisic acid responses early in development ^30^. Expression is strongly time-of-day dependent and exhibits some temperature dependence. The transcript peaks at lights-on, dropping quickly to a mid-day trough, and increasing again after dark, with higher peak expression at 20°C. HMCs 4, 9, 11, and 13 map to cell clusters of unknown state or type (Fig. 4A-D)^15^. Genes in HMC14 are expressed in two cell clusters, 3 and 15, which are identified as epidermal/trichome/trichoblast and vasculature, respectively (Fig. 4E)^15^. *ROOT AND POLLEN ARFGAP* (*RPA*), a gene reported to be involved in root hair cell growth via control of vesicle trafficking between the endoplasmic reticulum and Golgi apparatus, is expressed in this HMC, suggesting unstudied roles in these cell types^31^. *RPA* expression across the day is low and variable at 15°C. Expression is higher at 20°C, showing a slight oscillation increasing from ZT0 through ZT16, and dropping in the dark phase. HMC15 maps to vascular cluster CC15 (Fig. 4F). *CBL-INTERACTING PROTEIN KINASE 16* (*CIPK16*), which promotes tolerance to salinity, is expressed in this HMC^32^. Expression is not strongly temperature regulated, but it exhibits a strong oscillation, with peak expression at ZT0, dropping through the first half of the day to a trough at ZT8, and increasing again in the late night.

**Figure 4:**
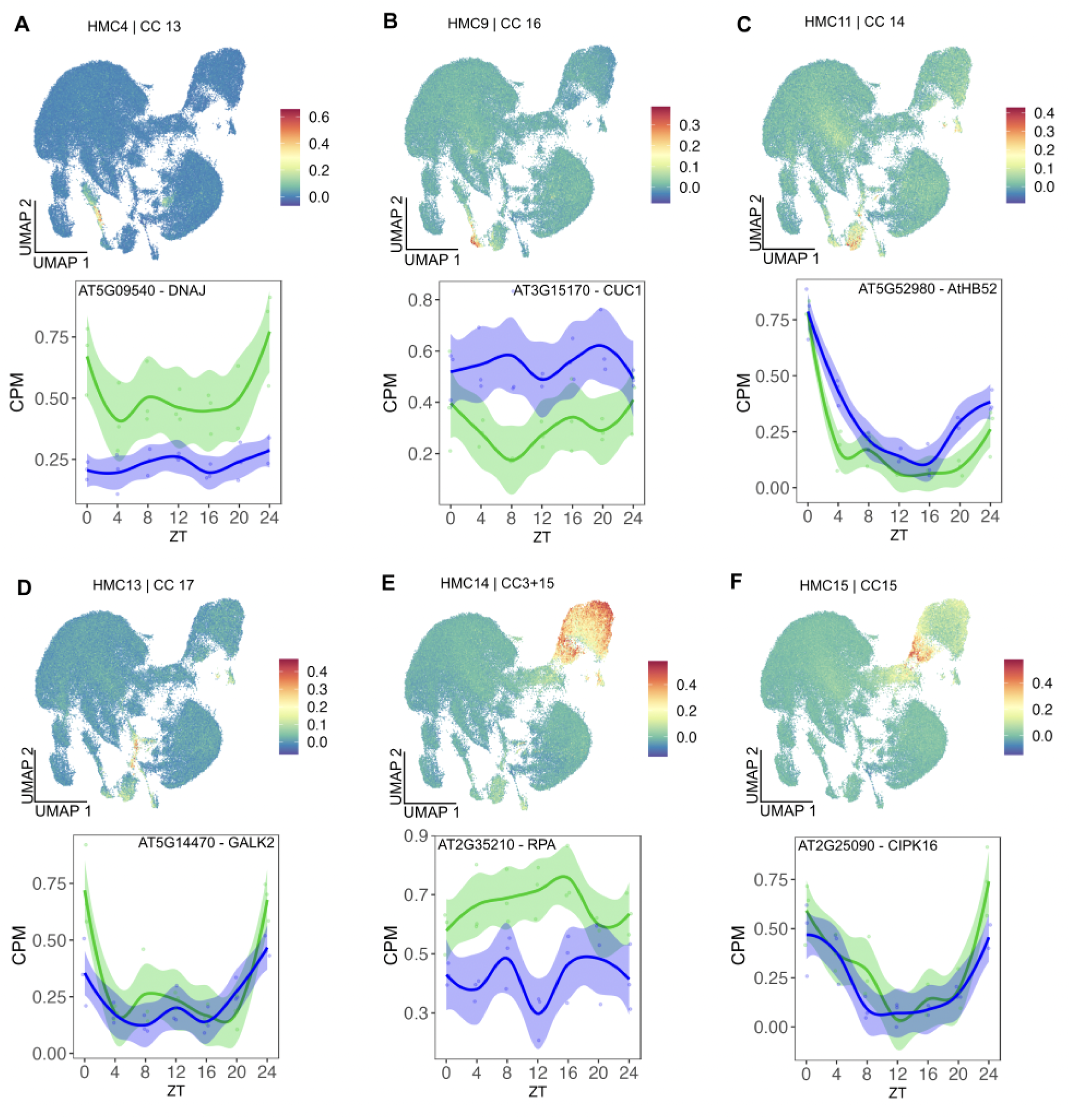
Tissue-specificity and time- and temperature-dependent expression of lowly-expressed genes. (A-F) UMAPs showing average expression of genes from selected clusters defined in the heatmap in Figure 3. HMC and CC (Cell Cluster) IDs are indicated above each UMAP. Color keys at the right of each panel indicate the range of expression. Time course expression of individual genes that were specifically retained by the mean CV<0.4 filter, included in each cluster, and determined to be differentially expressed under 15°C (blue) vs 20°C (green) in at least one time point are shown below each UMAP. Solid traces indicate the mean expression, and the shaded area surrounding the mean indicates the 95% confidence interval. Data from individual samples are shown as dots.

### Expression dynamics identify genes with shared biological processes within specific cell clusters

We previously used network-based co-expression analysis to identify genes regulated by PHOT2 and/or CAMTA2 under 15°C or 20°C^23^. Here we use this method to characterize temperature-dependent differences in Col-0, and map the resulting temperature-dependent communities to snRNAseq data. We show five examples of co-expressed gene communities that map to specific cell clusters (Fig. 5). Community 14 is characterized by increased expression at 15°C vs 20°C and corresponds to a single uncharacterized cell cluster (CC9)(Fig. 5A). This community shows specific enrichment for GO categories related to the cell cycle. Community 22 shows biphasic expression at 20°C and elevated expression at 15°C, peaking in the morning that maps to CC9 and CC10 (Fig. 5B). This community is specifically enriched for components involved in protein translation. Community 17 shows higher expression at 20°C than 15°C, and maps to CC11, which is identified as guard cells (Fig. 5C). This community is enriched for components of cell wall biogenesis. Community 25 shows sinusoidal expression with a peak at ZT8 and trough between ZT16-20 (Fig. 5D). Cycling amplitude of these genes is, on average, higher at 20°C. These genes mark CC11, and are enriched for core components of photosynthetic light-harvesting complexes. Community 6 is similar to Community 25, with peak expression at ZT12 at 20°C with a possible slight phase advance at 15°C, and also maps to CC11 (Fig. 5E). This community is enriched for starch and carbohydrate metabolism. The shared expression of communities 6 and 25 in CC11 is consistent with enhanced photosynthetic carbon fixation in these cells.

**Figure 5:**
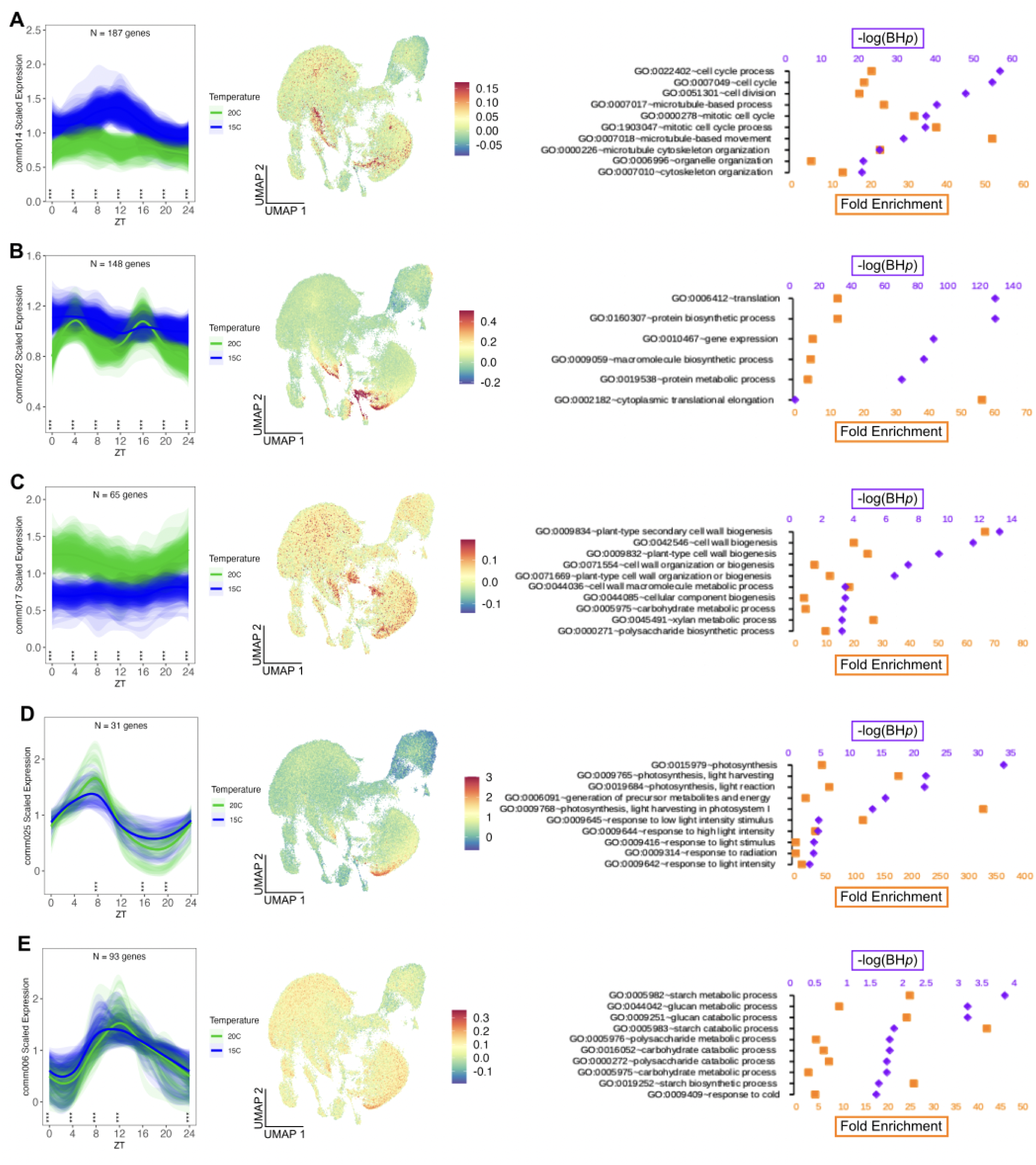
Gene communities defined by expression dynamics identify cell states or types. From left to right, Scaled gene expression, UMAP plots, and Gene Ontology enrichment are plotted for five communities of co-expressed genes (A-E). Communities were defined by scaled gene expression similarity (Pearson correlation >0.97). Average expression of the genes in the community are plotted in the UMAPs as in Figure 3. Black arrowheads indicate cell clusters of interest in the snRNAseq data set. The top 10 Gene Ontology (Biological Process) categories are shown at right. If fewer than 10 categories were enriched, all categories are shown. (A) Community 14 is expressed in an unidentified cell cluster in the UMAP. GO shows enrichment for Cell Cycle related genes. (B) Genes in community 22 mark the same cell clusters as (A), with some additional clusters. (C) Community 17 shows strong temperature-dependent differences in expression, and marks guard cell clusters in the UMAP. GO shows enrichment for cell wall components. (D) Community 25 s strong oscillatory gene expression peaking between ZT4-8. This community strongly marks an unidentified cell cluster in the UMA, along with high baseline expression across most regions of the UMAP. GO shows enrichment for core components of photosynthetic apparatus. (E) Community 6 genes are oscillatory, with peak expression between ZT8-12, with highest expression in the same cluster as comm25. GO shows enrichment for categories related to carbohydrate metabolism. (A-C) UMAPs were plotted including the argument ’max.cutoff = “q99”’ to optimize the visibility of specified cell groups.

We examined the phase of peak gene expression among the genes included in the co-expression networks generated from the CPM and mCV filters (Supp. Fig. 2C,D). Consistent with the result from the full transcriptomes, this subset of genes did not show large phase changes between temperatures. We therefore conclude that large-scale differences in the phase of gene expression oscillations are unlikely to be major factors in the temperature differences in physiology or development.

Taken together, we present analysis supporting the hypothesis that meaningful gene expression information can be retained by modifying the filtration strategy in standard RNAseq pipelines. This filtration strategy selects for higher-precision sequence information input to differential expression calculations, while increasing the proportion of identified differentially expressed genes. Combining the retained data from lowly-expressed genes with single-cell RNAseq data reveals clear specificity in expression patterns at the cellular level, and generates hypotheses regarding the tissue-specific roles of known developmental and environmental regulators, as well as uncharacterized genes expressed in defined cell types or states. We also show that while selection based on cell-type expression does not strongly enrich for specific pathways or processes, gene communities based on gene expression dynamics identify highly specific processes and cell clusters.

## DISCUSSION

### Filtering by the coefficient of variation of replicates increases discovery in RNAseq

Analysis of RNAseq data has become a common practice in biology, with well-established protocols and computational tools. Many of these protocols emphasize the need to filter out lowly-expressed genes due to measurement noise or unreliability^1,3–6,8^. While noise is undoubtedly a concern for low-level measurements (Supp. Fig.1A), the information discarded by pure expression level-based filtering may be biologically meaningful. We have not encountered published examples of filtering strategies based on the consistency among replicates. Here we present a method that directly addresses noise without sacrificing information due to low level expression. We found that filtering genes based on the coefficient of variation (CV) of the replicates retains genes with consistent measurements regardless of expression level, and preserves information about genes that may be expressed in few cells. Limited expression patterns may result from ensuring tight control over specific or sensitive processes. We find that transcription factors (TFs) are enriched among the genes specifically retained by the mCV filter, demonstrating the value of this new filter strategy (Fig. 2D,E). TFs are considered to be central factors in the regulation of physiology and development, with hierarchical gene regulatory networks of TFs controlling the expression of enzymes, signaling factors, and developmental determinants^33,34^. TFs are often not required at high intracellular concentrations, and as such can be expressed at comparatively low levels to carry out their functions^35^. Indeed, development and physiology may depend on tightly controlling TF expression^36,37^. Thus, the retention of TF information in RNAseq data is an important consideration for understanding cell- and tissue-specific processes.

Our filtration and analysis strategy is easily executed and generalizable to any RNAseq dataset that has at least three replicates per sample condition. RNAseq has become a ubiquitous strategy for the identification of transcriptional responses across all areas of biology. Each analysis pipeline includes a step for removing the least-informative genes from analysis^1,8,38^. This filtering step limits analysis to genes likely to yield biologically meaningful results. We show that by replacing the typical expression-level-based filter with one based on CV across replicate samples, we can recover data on biologically important genes expressed at very low levels. While we examine genes that respond to ambient temperature and the diurnal cycle in this study, RNAseq analysis pipelines are system-agnostic. As such, any transcriptomic study that includes sufficient replication can use this method. The RNAseq data used here were acquired at unusual read depth per sample (Fig. 2A). While this undoubtedly provides more information on lowly-expressed genes than experiments using more typical read depths, information on consistent gene expression that an expression-level filter woukd remove will be retained regardless of read depth^4,39,40^.

### Linking RNAseq with snRNAseq data to understand the cellular basis of environment-dependent gene expression

We preform two separate analyses to derive cell-specific information from genes identified in the RNAseq data. First, we specifically analyzed the cell-specific expression profiles of lowly expressed genes and found that their expression corresponded to specific cell clusters (Figs. 3-4). Second, we used network-based co-expression to uncover groups of genes with shared expression dynamics across the day, and found that some of these groups marked specific cell types or states (Fig. 5). Interestingly, we find that lowly expressed gene groups with enrichment in particular cells did not predict expression dynamics, with many wave forms represented across the day (Supp. Fig. 2). We conclude that cells themselves do not define expression dynamics and, consistent with previous studies, many temporal expression profiles can exist in a single cell^21^. However, we find that when temporal expression dynamics link genes, they may well occur in the same cells. Two notable co-expressed gene communities identified partially overlapping cell clusters that represent likely cell states, rather than cell types or fates (Fig. 5A,B). These gene communities were highly enriched in cell cycle and translation components, and the identity of the cell cluster was unknown^15^. Interestingly, we find that a gene characterized as a novel marker of proliferating cells in root and shoot meristems, *MERISTEM CELL CYCLE 1* (*MERCY1-*AT5G16250), is included in Comm014^41^. We suspect that other such markers are present in this community. The fact that cell cycle and translation components overlap within this cluster suggests that these cells are in a transitional state of growth and division, and may in fact represent multiple cell fates. Interestingly, HMC7 defined from the heatmap also shows enrichment in this cluster, and the genes in this group of lowly-expressed genes show a trend toward higher expression at 15°C than at 20°C, similar to the cell cycle regulators identified by co-expression (Supp. Fig. 2G, Fig. 5A). The genes in HMC7 are not enriched for specific GO categories, and the genes in HMC7 are not included in Comm014. HMC7 may represent multiple cell types that are simultaneously undergoing cell division.

The ability to capture carbon through photosynthesis is a defining feature of most plants. We found that co-expressed genes comprising the core photosynthetic light-harvesting complexes (Comm025) and starch metabolism (Comm006) show high expression in a single cell cluster (CC10) (Fig. 5D,E). CC10 is also highly enriched for translation (Fig. 5B). Photosynthesis genes were expressed with a peak at ∼ZT8 and higher-amplitude oscillations at warmer temperatures. Starch metabolism genes were expressed with a peak at ∼ZT12 at 20°C and ∼ZT8 at 15°C. While expression of these genes is not unique to these cells, the observed enrichment suggests that there is in fact a population of highly photosynthetic cells in the plant. This cell cluster also shows highly active translation, consistent with rapid production of these protein components (Fig. 5B). Consistent with this, the circadian time-course snRNAseq study found a range of cell groups enriched for photosystem I components, suggesting that these genes may be more enriched in particular cell types^21^. These components were enriched at Circadian Time (CT) 0-4, consistent with their morning expression. Even small modifications to photosynthesis lead to significant increases in yield^42^. Engineering the biochemistry of light harvesting, electron transport, and carbon capture is an active area of research, including potential engineering of cell-specific functions, as in C4 plants^43–47^. Consistent with previous studies, we found a highly photosynthetic cell population in *Arabidopsis*, a C3 plant, demonstrating that photosynthesis is not uniform across plant tissues^48,49^. Understanding the mechanisms underlying this variation may be important for increasing photosynthetic carbon capture and crop yield.

In sum, we demonstrate an improved CV-based strategy for filtering RNAseq data that retains high quality information lost with standard expression level filtering. This method is simple to incorporate into standard analysis pipelines and is broadly applicable to transcriptomic analysis across organisms. We provide proof-of-principle for our study, sowing that verified, biologically meaningful target genes are retained for analysis and localized to specific cell groups of unknown fate or function. We propose that this strategy can be used to mine existing datasets. As more data are reported for lowly-expressed genes, it may promote the development of increasingly detailed models of tissue-specific gene expression and gene regulatory networks.

## METHODS

### Coefficient of variation-based filter for RNA sequencing data

Our previously published gene expression data from *Arabidopsis* Col-0 plants grown under 20°C and 15°C and sampled at seven timepoints across day 10 [GEO accession GSE240117] were re-analyzed^23^. Gene-level reads were normalized to library size (counts per million reads, CPM) in edgeR^2^, accounting for all genes with non-zero values. The coefficient of variation (CV) was calculated from the CPM values for each biological replicate set (three replicates for each temperature and time point). The mean CV across all samples was calculated and used as the filtering criterion. Genes retained utilizing a range of CV thresholds were counted. Gene lists for thresholds of interest were retained. These lists were used to select gene-level read counts which were normalized to the new library size to obtain CPM values for further analysis. Downstream analysis was carried out using genes with a mean CV <0.4 (referred to in the text as mCV) unless otherwise noted. Differential expression was calculated in edgeR, with False Discovery Rate (FDR) ≤0.05 and Fold Change (FC)=|0.25|^23^. Filtering based on expression level was carried out as previously reported: retained genes had at least one sample with CPM>1 (defined by ’rowSums(cpm(TotalRawCounts_Col0.dge)>1) >=1’).

### Gene Ontology

Gene Ontology (GO) enrichment analysis was carried out in DAVID (https://david.ncifcrf.gov/home.jsp) as previously described, with an inclusion threshold of Benjamini-Hochberg *p*-value<0.05^23,50^.

### Mapping lowly-expressed genes within single nucleus RNAseq data

Lowly expressed genes were defined as genes with a summed expression <42 CPM across all 42 Col-0 samples (CPM<1 per sample) in our time course RNAseq data. We mapped these data to our published single-nucleus RNAseq data from 12-day-old seedlings [GEO accession GSE226097]^15^. Count data for these genes were extracted from the ‘GSE226097_seedling_12d_230221.rds.gz’ object and average expression per cell cluster was calculated in Seurat (version 4.3.0)^51^. Data were scaled per gene, and similarly expressed genes were identified using k-means clustering using ComplexHeatmap (version 2.13.1)^52^. Mean scaled expression of the genes in these heatmap clusters (referred to as HMC) was plotted per cell within the UMAP using Seurat. Expression values for individual genes (CPM) or gene groups (scaled CPM) were plotted using ggplot2 (version 3.4.3)^53^. Some UMAPs were generated including the argument ’max.cutoff = “q99”’ to limit the effect of the most lightly expressing cells on the lookup table and optimize the visibility of specified cell groups. These instances are noted in the figure legends.

### Network-based co-expression determination

Genes sharing gene expression dynamics by temperature and across the day were identified as previously described^23^. Briefly, differential expression between temperatures was calculated within the mCV transcriptome using edgeR, and genes showing a significant difference at any time point were retained. CPM values were scaled, and Pearson Correlation Coefficients were calculated and used to build a co-expression network. Co-expressed gene communities were identified using igraph (version 1.4.2)^54^. Mean expression for the gene group was plotted using ggplot2 and statistical differences were Resulting gene communities were then mapped to the day 12 snRNAseq UMAP using Seurat as above.

### Analysis of Time-of-Day Gene Expression

Samples were analyzed for time-of-day–dependent (cycling) expression as previously described^55^. Briefly, time-course expression data were formatted and processed using the HAYSTACK model-based framework for rhythmicity detection^20^. HAYSTACK operates by comparing observed expression profiles to a defined set of rhythmic models with known period, phase, and amplitude. Genes with a model correlation coefficient (R > 0.8) were classified as high-confidence cycling transcripts. The threshold for high-confidence phase differences was determined by the product of R values at each temperature (R_15°C_ x R_20°C_ >0.64).

## DATA AVAILABILITY

Data supporting the paper’s conclusions are included in the figures and supplementary data. The data sets analyzed in this study were previously published and are available via the Gene Expression Omnibus. Time-course RNAseq: GSE240117 ; snRNAseq: GSE226097.

## ACKNOWLEDGEMENTS

We thank Dr. Daphne Ezer (University of York) for helpful conversations. J.C. and J.R.E. are Investigators for the Howard Hughes Medical Institute.

## AUTHOR CONTRIBUTIONS

A.S. conceived and designed the study and analyzed data. T.A.L. designed experiments and analyzed data. N.T.H. and T.P.M. analyzed data. J.R.E. supervised research and acquired funding. J.C. supervised research and acquired funding. A.S. wrote the paper with input from the other authors.

## FIGURE LEGENDS

**Supplementary Figure 1:**
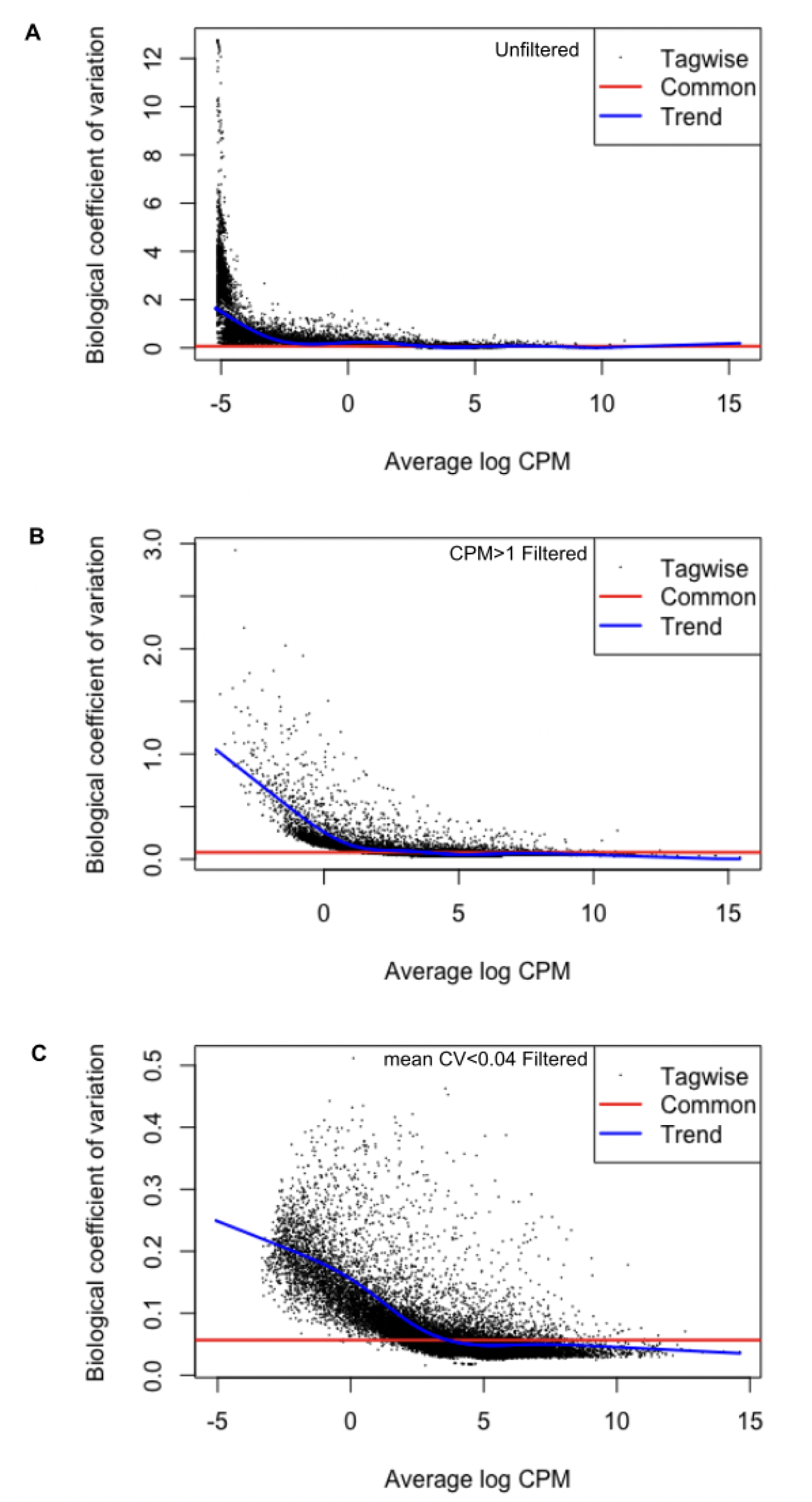
Biological coefficient of variation (BCV) by filter. Biological coefficient of variation by gene in (A) Unfiltered data, (B) CPM filtered data, and (C) mCV filtered data. Note the y-axes of the plots to compare the effects on the noise level of genes retained under each condition.

**Supplementary Figure 2:**
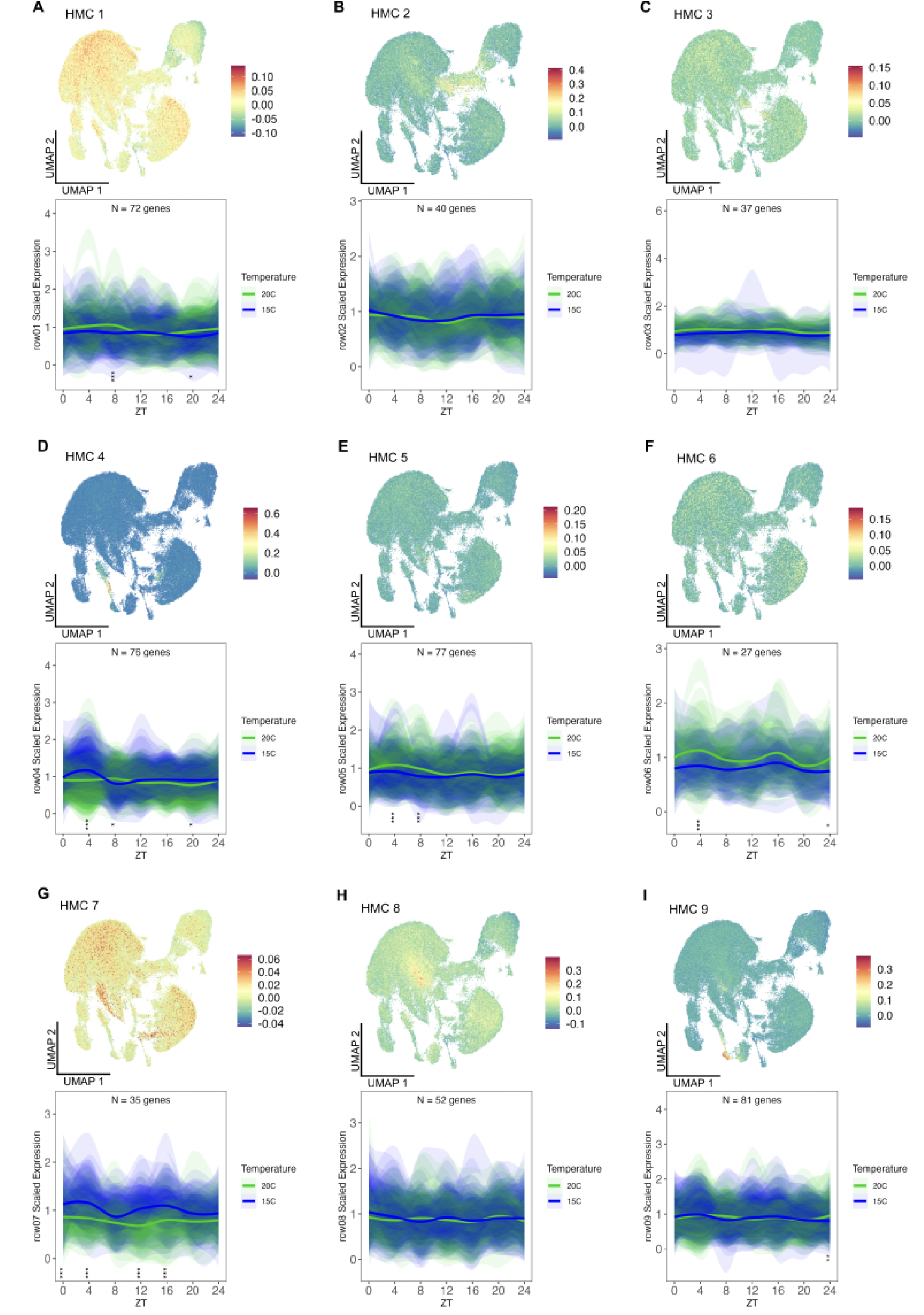
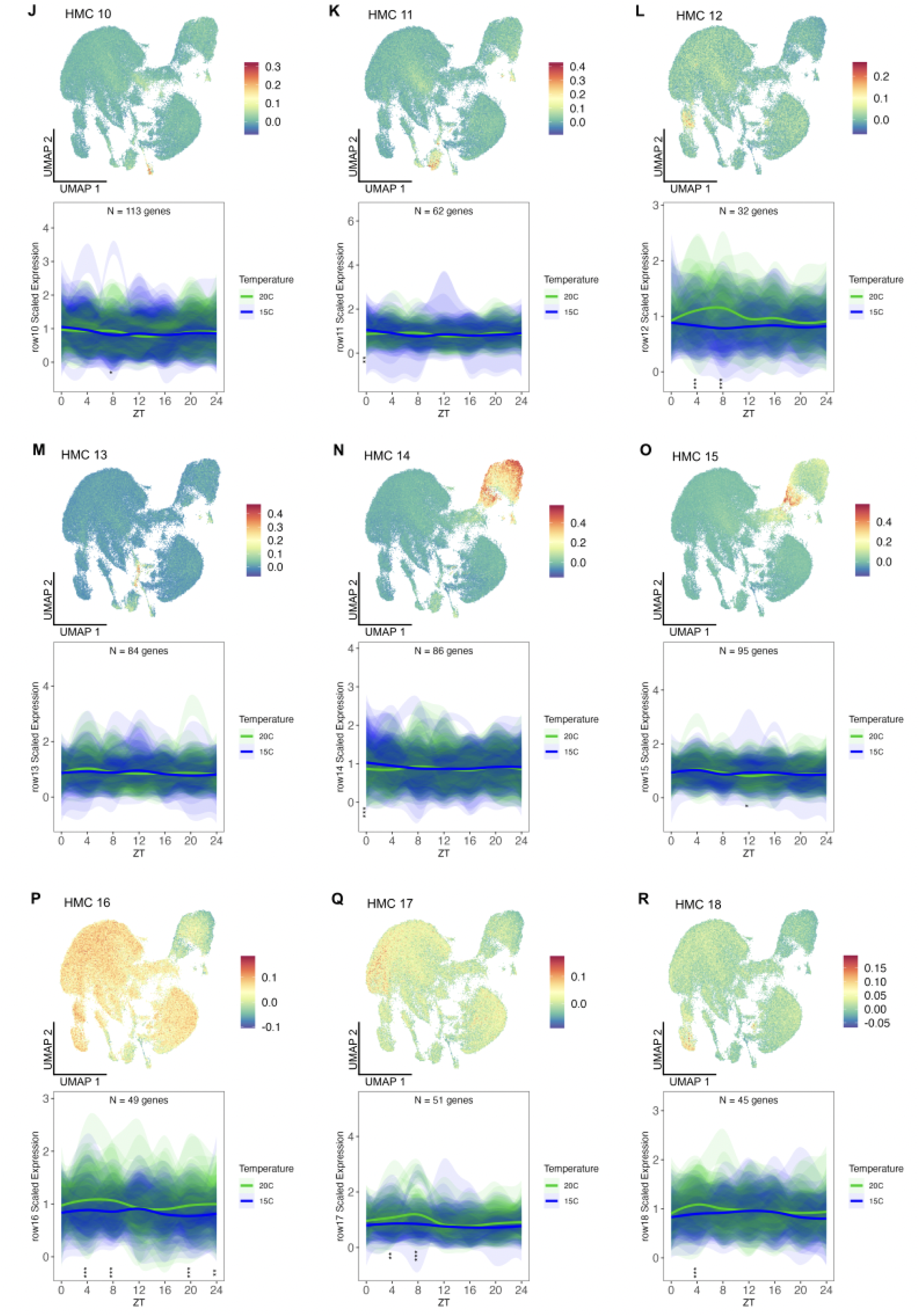
UMAP and time course gene expression plots for lowly expressed DEGs in each HMC. (A-R) UMAPs were generated based on the HMCs in Figure 3A. Scaled gene expression profiles for each HMC are shown below the respective UMAP, and colored by temperature (15°C - blue, 20°C - green). solid trace indicates the mean scaled expression of all genes, and shaded areas indicate the 95% confidence interval of each individual gene in the HMC. Asterisks near the x-axis indicate statistical significant differences between temperatures by two-way ANOVA (Time X Temperature) + Tukey’s HSD test (***0.001, **0.01, *0.05). The UMAP in (G) was plotted including the argument ’max.cutoff = “q99”’ to optimize the visibility of the cell groups.

**Supplementary Figure 3:**
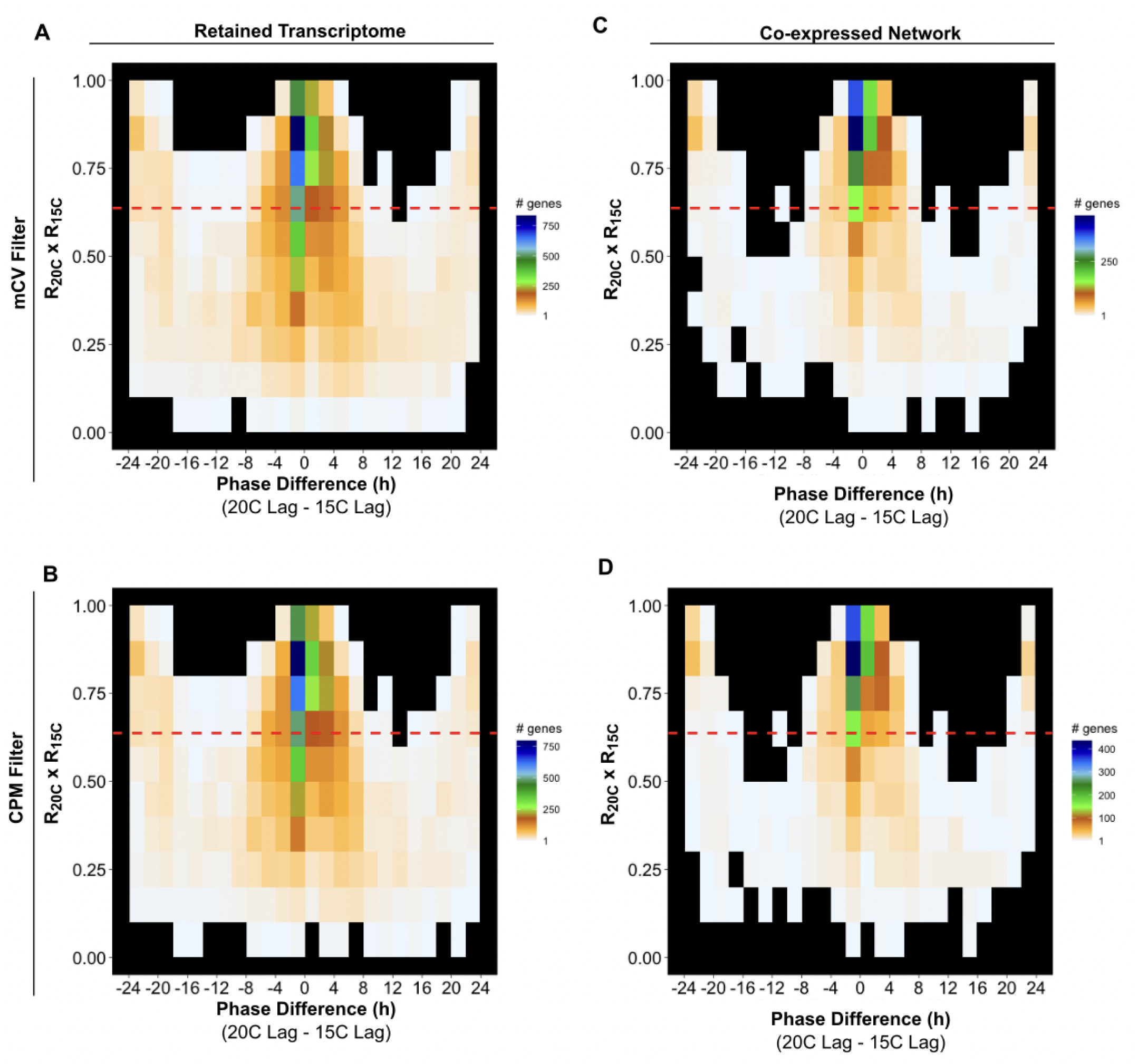
Minor phase differences between 15°C and 20°C. Diagrams show the distribution of phase differences in hours between 15°C and 20°C and the confidence measurement associated with the phase measurements. (A,B) Phase diagrams for the complete retained transcriptomes using the (A) mCV filter and (B) CPM filter. (C,D) Phase diagram of the genes included in the co-expression networks built from the (C) mCV filter and (D) CPM filter. Color lookup tables for gene density are shown to the right of each diagram. Red dashed lines indicate the threshold for high confidence phase measurements. For single genes, 0.8 is a typical confidence threshold for a reliable measurement. Here, the product of the two confidence measurements serves as the confidence level of the phase difference, thus leading to a threshold of 0.8 x 0.8 =0.64.

